# Massive intron gain in the most intron-rich eukaryotes is driven by introner-like transposable elements of unprecedented diversity and flexibility

**DOI:** 10.1101/2020.10.14.339549

**Authors:** Scott William Roy, Landen Gozashti, Bradley A. Bowser, Brooke N. Weinstein, Graham E. Larue

## Abstract

Spliceosomal introns, which interrupt nuclear genes and are removed from RNA transcripts by machinery termed spliceosomes, are ubiquitous features of eukaryotic nuclear genes [1]. Patterns of spliceosomal intron evolution are complex, with some lineages exhibiting virtually no intron creation while others experience thousands of intron gains [2–5]. One possibility is that this punctate phylogenetic distribution is explained by intron creation by Introner-Like Elements (ILEs), transposable elements capable of creating introns, with only those lineages harboring ILEs undergoing massive intron gain [6–10]. However, ILEs have been reported in only four lineages. Here we study intron evolution in dinoflagellates. The remarkable fragmentation of nuclear genes by spliceosomal introns reaches its apex in dinoflagellates, which have some twenty introns per gene [11,12]. Despite this, almost nothing is known about the molecular and evolutionary mechanisms governing dinoflagellate intron evolution. We reconstructed intron evolution in five dinoflagellate genomes, revealing a dynamic history of intron loss and gain. ILEs are found in 4/5 studied species. In one species, *Polarella glacialis*, we find an unprecedented diversity of ILEs, with ILE insertion leading to creation of some 12,253 introns, and with 15 separate families of ILEs accounting for at least 100 introns each. These ILE families range in mobilization mechanism, mechanism of intron creation, and flexibility of mechanism of intron creation. Comparison within and between ILE families provides evidence that biases in so-called intron phase, the distribution of introns relative to codon periodicity, are driven by ILE insertion site requirements [9,13,14]. Finally, we find evidence for multiple additional transformations of the spliceosomal system in dinoflagellates, including widespread loss of ancestral introns, and alterations in required, tolerated and favored splice motifs. These results reveal unappreciated intron creating elements diversity and spliceosomal evolutionary capacity, and suggest complex evolutionary dependencies shaping genome structures.

## Results and Discussion

### Ongoing intron gains in Symbiodinium species exhibit signatures of Introner-Like Elements

We reconstructed intron loss and gain in 562 sets of orthologs from five diverse dinoflagellate species using Malin [4,15], revealing a dynamic history of intron gain. Consistent with the high intron density of dinoflagellates, when scaled to rates of synonymous site turnover, rates of intron gain are far higher than previously reported for most other eukaryotes [2–5], on the order of one intron gain per kilobase per unit of synonymous change (i.e., dS) (dark blue and gray text in Figure 1a). Scrutiny of *Symbiodinium* introns revealed signatures of intron creation by DNA transposable elements (so-called -Like Elements or ILEs), with inverted repeats terminating at the 3’ splice site flanked by four nucleotide direct repeats, one of which serves as the 5’ splice site (Figure 1b,c) [6,9,10]. Allowing for near matches, this model accounted for between 17% and 28% of sets of candidate recent intron gains (Figure 1b). As expected, ILE sequence signatures were more pronounced for more recently gained introns, although signatures were also observed for introns shared across species, suggesting ILE-based intron gain occurring over long evolutionary time periods. BLAST searches revealed multiple families of ILE-derived introns for 3 out of 4 *Symbiodinium* species (Figure 1c).

**Figure 1.**
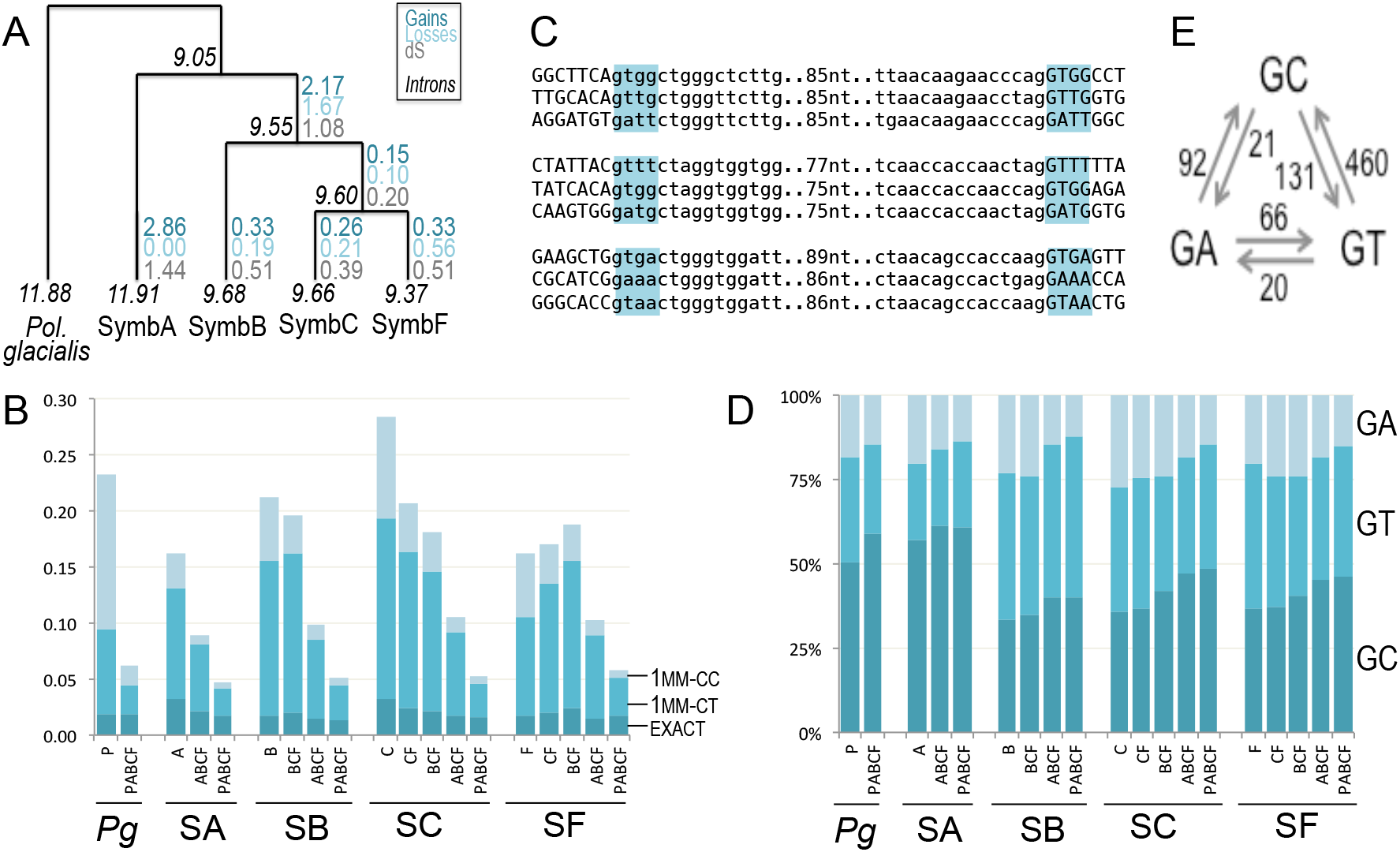
Intron-Like Element-Driven Evolution in Dinoflagellates. A. Reconstruction of intron density (italics) and intron loss and gain (dark/light blue) per kilobase. Rates of gains per intron gain per kilobase are of the same magnitude as dS (gray), contributing to increased intron density along all branches except the branch leading to SymbF. B. Substantial frequencies of dinoflagellate introns show signatures of gain by ILE insertion, with either a perfectly repeated tetramer and CT at nucleotides +5/+6 (dark blue), one mismatch from this pattern (medium blue), or one mismatch but with CC at +5/+6 (light blue). Introns are separated by estimated time of intron gain based on phylogenetic distribution of the position, with most recently gained introns to the left in each group. For instance, for *Symbiodinium* F (‘SF’), introns range from species specific (‘F’) to those shared with subsets of *Symbiodinium* species (CF/BCF/ABCF) to shared with *P. glacialis* (PABCF). C. Examples of class of ILEs for *Symbiodinium* clades A (top), B (middle) and F (bottom), showing 4 nucleotide target site duplication (TSD, blue). D. Boundary site usage, showing that GA (light blue) is more common for recently-gained introns (left, in each group) and GC (dark blue) is more commonly used in older introns (right). E. Identified boundary site conversions between *Symbiodinium* C+F species, showing biased conversion away from GA and towards GC.

Whereas nearly all spliceosomal introns across eukaryotes have a ‘GT’ 5’ splice site, previous work revealed high frequencies of GA and GC splice sites in dinoflagellates [11,12]. Indeed, we found high frequencies of atypical 5’ splice site variants (GA and GC) in all species, suggesting an early origin of splice site diversity within dinoflagellates. However the evolutionary history of GA and GC boundaries appears to be quite different. Both the greater frequency of GA among recently gained introns (Figure 1d) and a conversion bias from GA to other boundaries over evolutionary time (Figure 1e) are consistent with GA boundaries being suboptimal variants largely attributable to newly-gained introns. By contrast, GC boundaries show the opposite pattern, with overrepresentation among older intron classes and conversion bias towards GC boundaries over evolutionary time, suggesting that GC boundaries may be preferred over GA and even GT boundaries. Consistent with GC representing the optimal boundary, GC introns had statistically higher splicing efficiencies, although notably all classes had very high splicing efficiencies (median > 99%) within the available polyadenylated RNA-seq datasets (Supplemental Materials).

### Unprecedented diversity of ILE elements in P. glacialis

We next turned to the outgroup species, *P. glacialis*. Scrutiny of intron sequences revealed an abundance of ILE activity. Sequence similarity searches revealed 12,253 candidate ILE-derived introns, 92.5% (11,331) of which fell into 15 classes accounting for at least 100 introns each (Figure 2a; classes are numbered according to abundance). These classes exhibited a remarkable diversity of TSD lengths and strand flexibility, and flexibility of splicing pattern relative to insertion site, as we now describe (full sets of predicted ILEs are provided as Supplemental Dataset 1).

**Figure 2.**
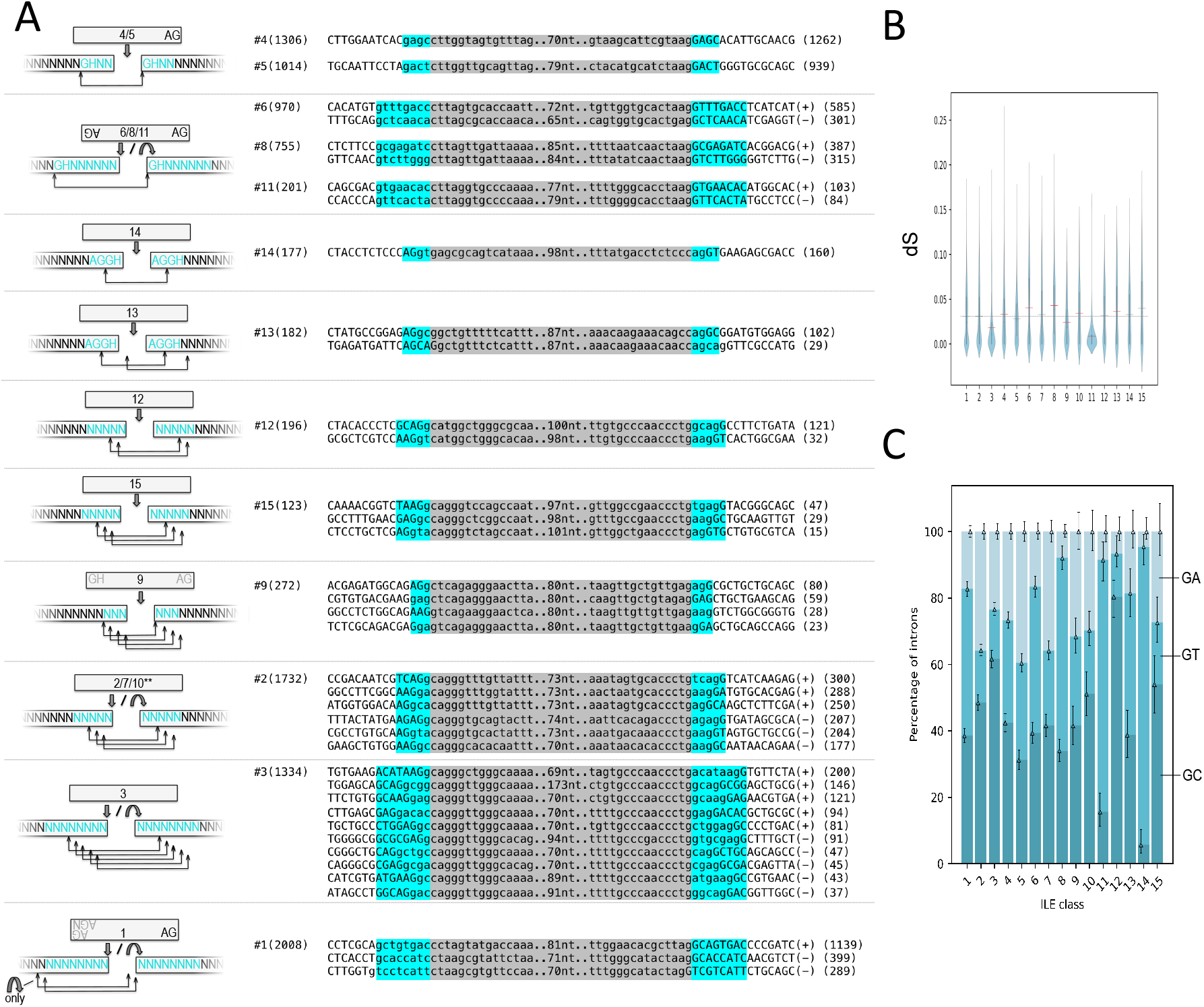
Diversity of Introner-Like Elements in *P. glacialis*. A. Diversity of ILEs. 15 classes of ILEs are shown. Diagrams at left of the mechanisms of insertion and intron creation are shown, indlcating class number(5) sequence of TSD (blue), strand orientations in which introns are created (single strand=straight arrow; both strands=straight/curved arrow), and positions of splicing boundaries (square bracket arrows below). Total number per class is shown in parentheses, followed by examples in various frames and strand orientations, with numbers in each orientation given in following parentheses. B. Difference in ages of ILEs across age classes. For each class, distributions of terminal branch lengths for ILE phylogenetic tree are shown. Red indicates significant deviation from the mean across classes (horizontal line). C. Difference in splice site boundary across classes, with light/medium/dark representing GA/GT/GC.

Among the 15 classes responsible for the creation of >100 introns, 5 classes showed a relatively simple pattern such as that previously reported in *Aureococcus anophagefferens*, in which one splice site is carried within the ILE and the other is recruited from the TSD upon insertion (Figure 2a). For instance, elements from classes 4 and 5 (1306 and 1014 introns, respectively) showed a *Symbiodinium*-like pattern, with a 4nt TSD with general sequence GHNN (where H=A/C/T), and with insertion leading to intron creation in only one orientation, and with a consistent positional relationship (with one end of the ILE serving as the 3’ splice site and the 5’ end of one TSD serving as the 5’ splice site). Classes 6, 8 and 10 (970, 755 and 201 introns) showed similar patterns but with a different cut site length (8 nts, GHN6), and with an ability to create introns when inserted in either orientation, in a pattern similar to that reported for *A. anophagefferens* [9] (interestingly, for all classes, there was a bias towards one orientation over the other). Class 14 showed a novel pattern, in which the 5’ and 3’ splice sites are both recruited from a 4 nt TSD of AGGH. Even among these relatively simple patterns, it is remarkable to find ILEs within the same species with different characteristic TSDs, as it indicates intron creation by different transposable element mobilization machineries.

Other classes revealed even more complex patterns, in which ILEs could be created in multiple frames. For instance, classes 13, 12, 15 and 9 (182,196,123 and 272 introns, respectively) showed the ability to create introns on a single strand orientation but in between 2 and 4 different frames. Class 12 (TSD=5nt) insertions consistently recruited the 3’ splice site from the TSD, but the splice site could be in multiple positions within the TSD (either N<u>AG</u>GH or NN<u>AG</u>G), with the 5’ splice site recruited either from the TSD (NAGGH) or from the terminal G of the TSD (NNAG<u>G</u>) along with the initial ‘C’ of the ILE. Class 13 (TSD=4nt) could either recruit both splice sites from the TSD (AG<u>GH</u>) or use the 2^nd^ and 3^rd^ nucleotides of the ILE as a 5’ splice site (G<u>GC</u>) while recruiting the 3’ splice site from the last nucleotide of a NNNA TSD and an immediately downstream G (i.e., <u>NNNA</u>|Ggc…nn|<u>nnna</u>gNNN, where “|” marks the ILE/TSD boundaries, underline indicates the TSD, and upper/lowercase indicates the exonic/intronic sequence). For class 9 (TSD=3nt), we observed insertions in four different frames: the 5’ splice site can fall within the TSD, within the ILE, or can span the ILE-TSD boundary, and the same is the case for the 3’ splice site. Classes 2, 7, 10 and 3. exhibited similar flexibility of frame, in addition to insertion in either orientation. Most remarkably, class 3 (1334 introns, TSD=8nts) created introns in 5 different frames by inserting in either strand orientation, for a total of 10 different mechanisms of intron creation for a single ILE class.

Finally, class 1 showed an interesting strand dependency. When inserted in the most common orientation, it showed a ‘simple’ pattern, with all introns using the 5’ end of the TSD and the 3’ end of the ILE as splice sites. By contrast, when inserted in the alternative strand orientation, because of diversity of the end of the ILE (either TAAG or TAGG), splice sites could derive from either the 5’ end of the TSD and the 3’ end of the ILE (as in the more common orientation), or from one nucleotide site upstream (5’ splice site from the first pre-TSD nucleotide plus the first TSD nucleotide; 3’ splice site from the 2nd- and 3rd--‐to-last nucleotides of the ILE).

Further scrutiny of the different ILE classes revealed differences in various other features. First, ILE classes differed significantly in their 5’ splice site usage (GT/GC/GA; Figure 2b). Some of these patterns were consistent with the mechanism of intron creation: for instance, class 12 ILE predominantly used GC 5’ splice sites, consistent with the initial ‘C’ base of the ILE serving as the second base of the intron. The reasons for other biases were not immediately clear (for instance, the strong preference for GT for the TSD-derived splice site for class 14). Second, we also observed differences in age distributions of ILEs of different classes, consistent with ongoing intron gain over a long period of time within dinoflagellate evolution (Figure 2c). For instance, in the phylogenetic tree connecting ILEs of class 11, the median terminal tip length in sequence divergence was 0.9%, compared to 3.8% for class 8.

### Recruitment of splice sites from coding sequence predicts intron phase bias

These data provide an unprecedented opportunity to understand the determinants of intron positions within genes. Of particular interest is the so-called phase bias, in which intron positions across diverse species tend to fall between codons (‘phase 0’) rather than after the first or second nucleotide of a codon (phase 1 and 2, respectively) [9,16,17]. Previous work has suggested that, because most characterized ILEs recruit splice sites from preexisting sequences (TSDs and/or flanking sequence), biases in the distribution of nucleotides and nucleotide motifs in coding sequences could explain phase bias [9,14,17]. Indeed, the distribution of flanking nucleotides strongly predicts the phase bias across ILE classes (*R*^2^ = 0.90, *P* < 10-^5^) (Figure 3a). A second prediction of the model has not been tested: if phase bias is due to recruitment of constrained splice sites from exonic nucleotides, then recruiting more splice site nucleotides from exonic sequence (TSD or flanking sequence) should lead to a stronger phase bias. For instance, we would expect stronger phase bias for class 14 ILEs, which recruit all four splice site nucleotides from a AGGH TSD, than for class 4 ILEs, which recruit only two from a GHNN TSD. This pattern is indeed observed: among ILE classes which recruit a consistent number of nucleotides, ILE classes that recruit more nucleotides show stronger phase bias (Figure 3b; *P* < 10^-4^ by randomization). An even more direct test takes advantage of ILE classes that vary in the number of recruited nucleotides based on the frame of insertion. As noted above, class 13 ILEs can create introns in two different ways. For some insertions, all four splice site nucleotides are recruited from a AGGH exonic TSD, whereas for others, only the 3’ splice site is recruited from exonic sequence. As expected, class 13 ILEs that recruited four splice site nucleotides show stronger phase bias (63.0%:23.6%:13.4% phase 0:1:2) than do class 13 ILEs that recruited two splice site nucleotides (51.1%:23.4%:23.5%). Among eight cases in which a single intron class differed by insertion frame in the number of coding nucleotides recruited, we found a correspondence in the expected direction in seven (with no difference in the eighth), with both phase 0 bias and overall bias higher among insertions in the frame that recruited more nucleotides (Figure 3c; *P* = 0.008 by a binomial test). These results constitute the most direct test available of the so-called ‘proto-splice site’ model, in which biases in the sequences into which new introns are created strongly predicts the phase bias of the resulting introns [14,17].

**Figure 3.**
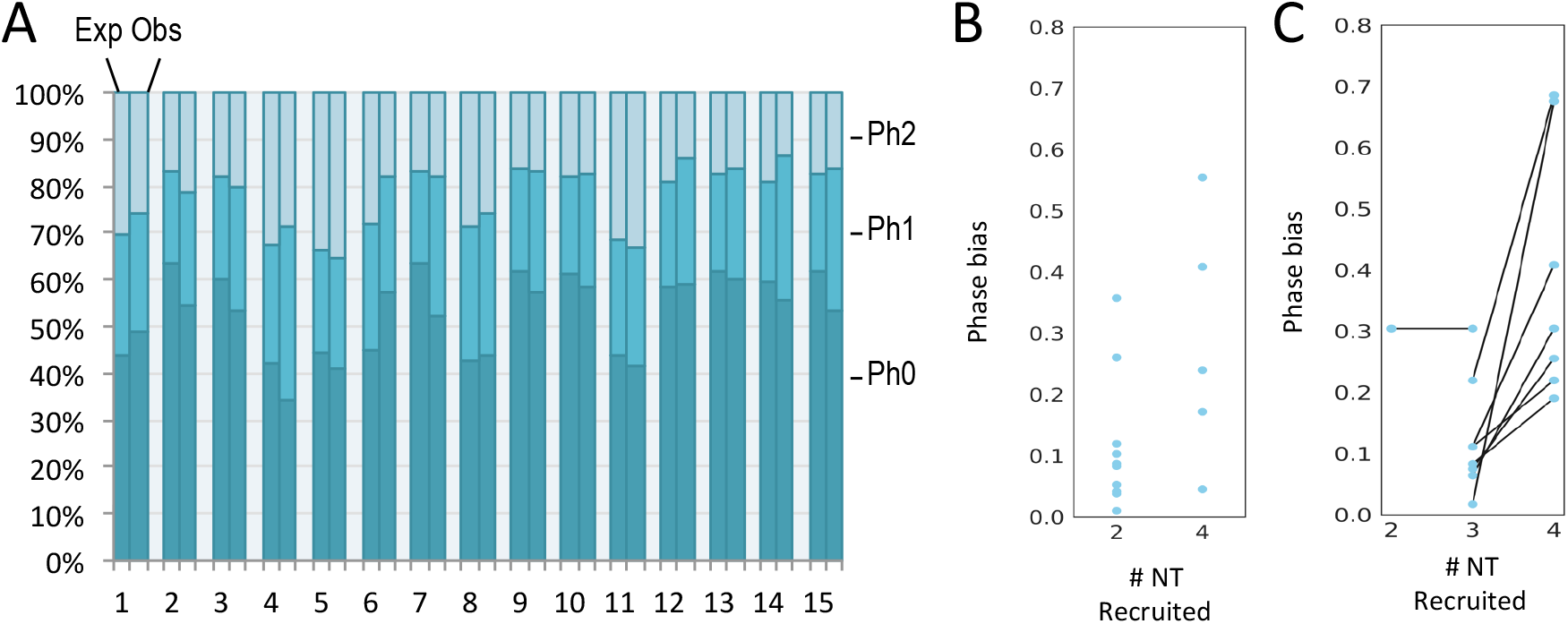
Intron phase bias in *P. glacialis* is predicted by recruitment of exonic nucleotides. A. Expected and observed phase distribution for ILEs in each class. Expected is based on the distribution relative to coding frame of the observed tetramers flanking introns. B. Phase bias for classes of ILEs based on the number of recruited exonic nucleotides, for classes that consistently recruit the same number of exonic nucleotides. Phase bias is calculated as the fractional excess of repeatability, that is 3(*p*_0_^2^+*p*_1_^2^+*p*_2_^2^)-1, where *p* values indicates the fraction of introns in each phase. C. Phase bias for classes of ILEs that have varying numbers of recruited exonic nucleotides depending on different frames of insertion, as a function of numbers of nucleotides recruited.

### Transformation of dinoflagellate spliceosomal systems

We also observed that dinoflagellate intron-exon structures appear transformed in various ways. Comparison with diverse eukaryotes revealed very few shared intron positions, indicating nearly complete intron loss by the time of the ancestor of the studied species (Figure 4a). Dinoflagellate splice sites are also transformed, with a weakened extended 5’ splice site (GHNNNN) but a highly conserved downstream exonic motif of GH directly downstream of the 3’ splice site, a pattern consistent with these dinucleotides’ relationship to the 5’ splice site (also GH) in ILE-created introns (Figure 4b). Interestingly scrutiny of the spliceosomal machinery did not reveal large-scale transformation (Figure 4c), in contrast to the spliceosomal simplification observed in some but not all intron--‐poor lineages [18–21].

**Figure 4.**
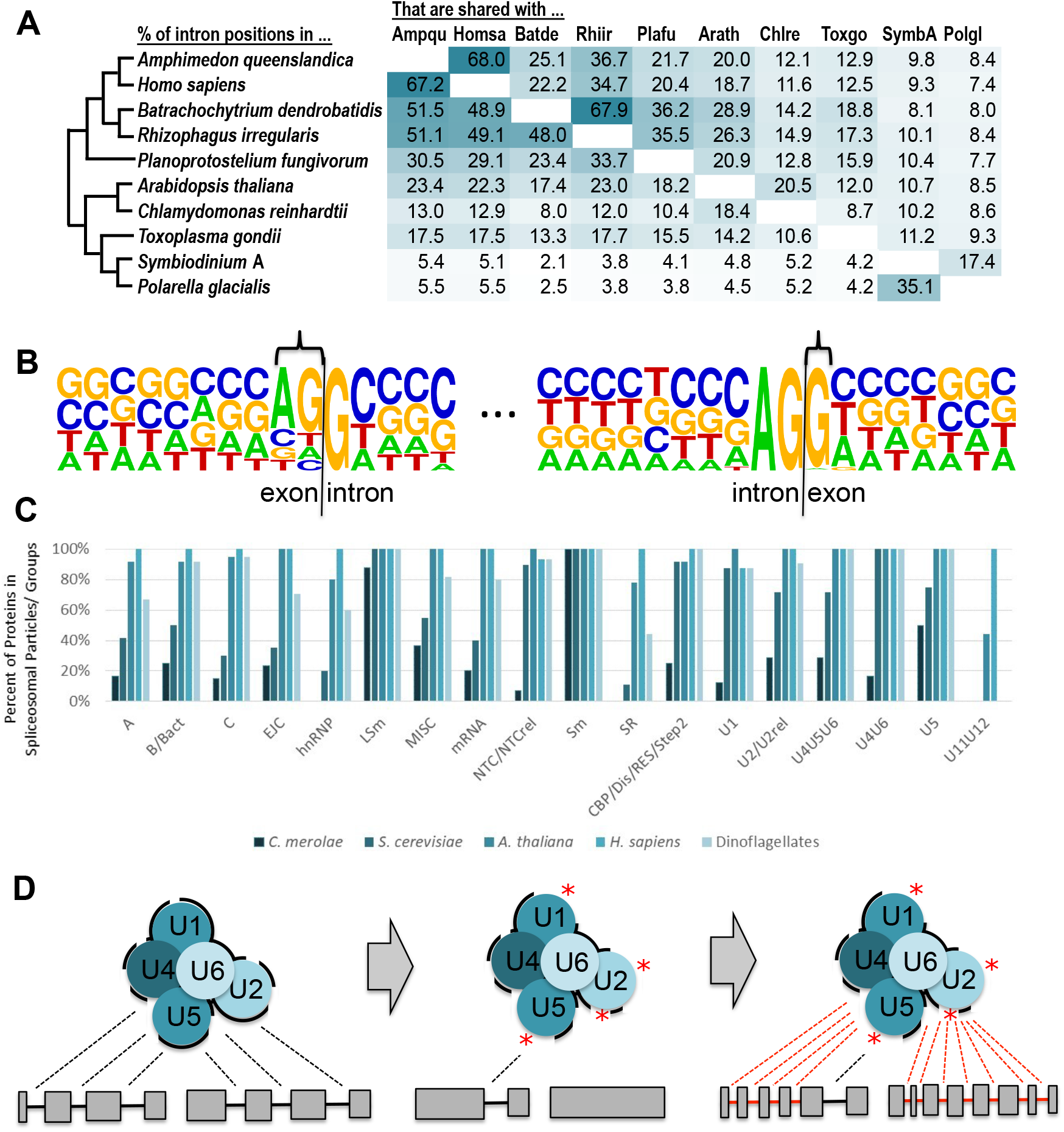
Transformation of the dinoflagellate spliceosomal system. A. The representative dinoflagellates *Symbiodinium* A and *P. glacialis* share few of the intron positions found in various eukaryotes, suggesting widespread intron loss. B. Consensus sequences from *P. glacialis*, showing increased conservation of exonic bases at the −2 and −1 and +1 positions (bracketed). C. Percent of putatively ancestral spliceosomal proteins retained for various spliceosomal complexes for five species, showing massive and moderate reduction in spliceosomal protein number for the red alga *C. merolae* and the yeast *S. cerevisiae*, respectively, but no evidence of substantial loss in dinoflagellates. D.Model for the evolutionary history of spliceosomal machineries in species with widespread intron gain. Ancestral spliceosomes are highly constrained (represented by thick black lines) by virtue of having to recognize large numbers of introns in intron---rich genes (box/line structures at bottom). Massive intron loss leads to reduced constraint (represented by red asterisks). Newly unconstrained regions allow for the recognition of newly-spreading ILEs (red introns).

### On complex path dependencies in the evolution of genome complexity

Our results reveal a highly transformed spliceosomal system in dinoflagellates, with ancient loss of ancestral intron complement and transformation of splicing signals coupled with ongoing and perhaps generally more recent intron proliferation. *Symbiodinium* thus joins the green alga *M. pusilla* [6] and the tunicate *Fritillaria borealis* [10] as lineages that couple three rare and partially opposing phenomena: massive loss of ancestral introns, transformation of splicing signals, and intron creation by ILE proliferation. This provides further support for the notion that the persistence of many ancestral introns may impose constraints on the spliceosome that reduce the capacity of the spliceosome to respond to invading transposable elements, reducing the probability of the emergence of ILEs (Figure 4d). This ‘law of unintended consequences’ wherein massive intron loss facilitates massive intron gain is strikingly opposed to the notion that some greater governing factor lends a given lineage a tendency for genome complexity to either increase or decrease, and underscores the complex dynamics governing the origins of genome complexity.

### Concluding remarks

By showing that the most intron dense known eukaryotic genomes have been created in substantial part through invasion of diverse Introner-Like Elements, these results establish ILEs as a force in creating substantial genome complexity in eukaryotes and support their role as major creators of introns. By revealing that dinoflagellates reached their unparalleled intron densities through massive intron loss coupled to the activity of an unprecedented diversity of intron-creating transposable elements, these results urge caution in the adoption of grand theories of genome evolution and instead suggest important roles for path dependency, stochasticity, and rates of genome architecture-altering mutations in the origins of genome architecture [22–25].

## Supplemental Methods

### Scripts and programs used

Scripts used for each analysis including data source download, with additional documentation, are available at https://github.com/Brookesloci/Simbiodinium_intron-er---_evolution, as are intermediate and final dataset files for the reported analyses.

### Data sources

We downloaded the genomes and annotations of *Symbiodinium goreaui* (clade C1) and *Symbiodinium kawagutii* (clade F) from http://symbs.reefgenomics.org/download/. We downloaded the genome and annotation of clade B from https://marinegenomics.oist.jp/symb/viewer/download?project_id=21. We downloaded the genome and annotation of *Polarella glacialis* CCMP1383 from the genome project site (https://espace.library.uq.edu.au/view/UQ:d71d29b). We downloaded the genomes and annotations of *S. microadriaticum* (GCA_001939145.1), *H. sapiens* (GRCh38)*, A. queenslandica* (GCA_000090795.1)*, B. dendrobatidis* (GCF_000203795.1)*, R. irregularis* (GCA_000439145.3)*, P. fungivorum* (GCA_003024175.1) *A. thaliana* (GCF_000001735.4)*, C. reinhardtii* (GCA_000002595.2) and *T. gondii* (GCF_000006565.2) from NCBI.

### Intron and HI analysis

Orthologs were defined by reciprocal blast hits. For multi-way ortholog sets, ortholog sets were defined conservatively, namely as a set with one gene per species, in which each gene in the set was defined as the pairwise ortholog for all other genes in the set. Ortholog pairs/sets were aligned at the protein level by standalone Clustal Omega 1.2.4, nucleotide sequences including intron positions and sequences mapped onto the resulting alignments. Confident alignment positions were defined as those for which there was 40% amino acid identity between each pair of orthologs over a window of 10 amino acid positions in each direction. dS was estimated using default parameters in PAML 4.9.

### ILE identification

ILEs were identified by all-against-all blastn searches with a cutoff of E-10, and with only pairs of introns where similarity (as defined by the boundaries of the blast hit) began and ended near both splice site boundaries (defined between the 5th base into the exon and the 10th base into the intron). Families of ILEs were defined using a greedy clustering algorithm.

### Reconstruction of intron loss and gain

We mapped homologous intron positions, discarding genes that lacked shared intron positions to avoid the inclusion of retroposed paralogs. The analysis was further restricted to include only intron positions flanked in the protein sequence alignment by four non-gap amino acid positions on both sides, leaving 6,832 intron sites to reconstruct intron gain and loss using a constant rate implementation of Malin, the probabilistic model of intron evolution developed by Csűrös *et al*. (Csurös, 2008). Rates of intron loss and gain were used to calculate a table of intron sites with a certain probability of presence, gain, or loss at individual intron sites for each of the four internal nodes, starting from the standard null model with 1000 optimization rounds and a likelihood convergence threshold of 0.001. Finally, a Markov Chain Monte Carlo (MCMC) method was employed to assess confidence by sampling model parameters and ancestral intron reconstructions by their posterior distributions for 100 replicates.

### Splicing efficiency analysis

For each species, splice junction sequences corresponding to the unspliced (5’ and 3’ exon-intron) and spliced (exon-exon) form of each annotated intron were constructed in silico for the longest isoform of each gene. Bowtie (v1.2.2) was used to map uniquely-mapping RNA-seq reads from each sample (using non-default parameter `-m 1` with all reads treated as unpaired) to the junction sequences, and reads overlapping each junction by ≥5 nt were used to calculate per-junction coverage values (adjusted to account for the number of mappable positions present in each junction construct). For each sample, introns with evidence of splicing (i.e. at least one read mapped across the exon-exon junction construct) and ≥10 reads mapped across one or more junctions were included in the splicing efficiency analysis. For such introns, we defined splice- and retention-supporting values as the exon-exon junction coverage and average of the two intronic-sequence-containing junction coverages, respectively. Overall splicing efficiency was then calculated by dividing the splice-supporting junction coverage (C_s_) by the sum of the retention-supporting junctions’ coverage (C_r_) and the splice-supporting junction coverage, i.e. Cs(Cr/2) + Cs100%. Lastly, each intron’s splicing efficiency was averaged across all RNA-seq samples where evidence of splicing of the intron was found to produce the final splicing efficiency data. Splicing efficiency values were compared between datasets through a Mann-Whitney test, showing significantly greater splicing efficiency for GC introns (*P* < 10^−5^).

### Spliceosomal protein searches

Spliceosomal protein searches were performed on proteome assemblies available from NCBI (*Saccharomyces cerevisiae*, R64; *Homo sapiens*, GRCh38.p12; and *Cyanidioschyzon merolae*, ASM9120v1) and TAIR (*Arabidopsis thaliana*, Araport11). Spliceosomal protein searches were performed as in Hudson et al. (REF). Briefly, sets of spliceosomal proteins were generated by using data from previous studies. To qualify for our search lists, the spliceosomal protein must be present in at least two of *H. sapiens, S. cerevisiae*, and *A. thaliana* proteomes (Fabrizio et al., 2009, Hegele et al., 2012, Koncz et a., 2012). Additionally, the Core snRNP and Non-core snRNP protein sets were defined from previous studies (Will and Luhrmann, 2011; Burge, 1999). Human and Arabidopsis spliceosomal proteins were used as queries in local BLASTp (version 2.9.0+) searches against independent *Symbiodinium* and *P. glacialis* proteome databases (initial e-value threshold of 10-6 (Atschul et al., 1997). The results from the BLAST searches were further screened by analyzing domain content (HMMsearch, HMMer 3.1b2 – default parameters), size comparisons against Human, yeast, and Arabidopsis (within 25% variation), and reciprocal best-hit BLAST searches (RBH) to the query proteome (Johnson et al., 2010; cite RBH). Ortholog candidates were accepted if they contained all the conserved domains, were within an acceptable size range. Candidates that did not meet the above criteria were manually screened. Initial analyses showed a large number of genes for which a subset but not all dinoflagellate species had a candidate gene. In some cases, a protein candidate was accepted even if it was outside of the size range as long as it still matched as an RBH as there are several examples of size transformations between spliceosomal orthologs in model organisms.. Given that dinoflagellate gene structures are highly atypical and thus presumably challenging to annotate, we performed BUSCO 3.0.1 (Simaeo et al., 2015) analyses of the five proteomes, which indeed revealed low scores (45.9% to 60.7% of Complete eukaryotic BUSCO groups). Therefore, the individual results from the *Symbiodinium* and *P. glacialis* searches were pooled, such that a gene found in a subset of the five species was called as ‘present.’ Results are shown in Supplemental Figure 1.

### ILE phylogeny analysis

We first removed the target site duplications present in each of the fifteen *Symbiodinium* ILEs to compare phylogenetic trees of only the ILE sequences. We inferred maximum likelihood (ML) phylogenetic trees using IQ---TREE v1.6.10 (Nguyen et al.,2015) and ModelFinder (Kalyaanamoorthy et al., 2017) to find the optimal model for each ILE. We then inferred the most likely tree for each again using IQ-TREE, with bootstrap scores inferred using the ultrafast bootstrap method (Hoang et al.,2018). Final trees were rooted by midpoint.

The ILE sequences are a problematic phylogenetic dataset because they are short with overall low signal, causing tree selection to fluctuate. Therefore, to increase confidence in our trees, we used the approximately unbiased (AU) test (Shimodaira, 2002) to check that the optimal tree was not designated optimal by chance. We invoked ten independent ML phylogenies for each ILE family (as above) and then performed the AU test with each ILE family set of ten trees. For 14 out of the 15 ILE families, all 10 trees were accepted with p-values > 0.05. However, ILE family 8 consistently had one tree out of 10 rejected. In this case, we next applied the more conservative test of Shimodaira and Hasegawa (1999), and all ten trees were accepted.

### Estimation of ILE age

For each ILE phylogeny, lengths of each tip branch of the tree were extracted as a proxy of the age of the insertions. These distributions were then analyzed using a Mann-Whitney test to identify families with an average tip length statistically greater or less than the average across families.

### Splice site conversions

To identify instances of splice site conversion, we identified shared intron sites between *Symbiodinium* species C and F in which both species shared evidence for the same 4nt TSD, that is, where both species’ 5’ intron splice sites and both species’ downstream exons shared the same 4nt motif, with the exception of one mismatched base (the candidate mutant base) at the +2 position of one of the two species’ 5’ intron splice site. For example, if species C’s sequence was NNNNgact…agGACT and species F’s sequence was NNNNgcct…agGACT, then a A→C mutation was inferred.

### Phase calculations

For each set of ILEs, expected phase distribution was calculated as follows. For each intron, the two upstream and two downstream bases were concatenated to yield a tetramer that represents the possible intron insertion site (that is, the exonic −2, −1, +1 and +2 bases). The frequencies of these tetramers were compiled across introns in the set. For each observed tetramer, the frequency of this tetramer in the three phases was calculated across all coding sequences from the same species. The phase distribution for each phase (0-2) was then estimated as the weighted frequency for that phase across all observed tetramer insertion sites. Phase bias was defined in terms of the excess repeatability of phase across introns, defined as the probability that two randomly chosen introns from the set have the same phase minus the probability that two randomly chosen introns would have the same phase under equal phase distribution (⅓), divided by the random expectation (⅓). That is, [(*p*_0_^2^+*p*_1_^2^+*p*_2_^2^)-⅓]/(⅓), or 3(*p*_0_^2^+*p*_1_^2^+*p*_2_^2^)-1.

